# Exposure to paired blue light and nutrient alters aversive responses in Physarum polycephalum

**DOI:** 10.64898/2026.09.09.750388

**Authors:** Qiuran Wang, Niels Gaspers, Tobias Schlicht, Liberty Severs

**Affiliations:** Institute for Philosophy II, Ruhr-University Bochum; Faculty of Sciences, University of Lisbon

**Keywords:** Physarum polycephalum, Pavlovian conditioning, associative learning, basal cognition

## Abstract

Whilst forms of non-associative learning such as habituation are widely reported in single-celled organisms, whether associative learning is possible in systems without a nervous system is contested. In this study, we implemented a conditioning paradigm in the acellular slime mould *Physarum polycephalum* through repeated pairings of nutrients with an aversive blue light stimulus. We first established that blue light is aversive under our conditioning protocol, slowing migration and producing broader, less directed growth, confirming its well documented effect on behaviour. Subjects were then trained in up to four training sessions by placing an oat stimulus inside a region with blue light before testing with the light presented alone. Trained subjects were more likely to reach the blue light region compared to untrained controls, with shorter latency, suggesting that *Physarum polycephalum* weakens its avoidance of blue light after the region has repeatedly coincided with a nutrient. However, engagement did not strengthen with additional pairings, a pattern that admits multiple competing explanations in which associative and non-associative processes plausibly contribute to more transient associative-like effects. We conclude that a history of co-presentation is sufficient to relax an aversive response in *Physarum* but cannot, within the present study, attribute that change to associative learning.

## 1. Introduction

Pavlovian conditioning is generally considered to be an exemplary case of associative learning, wherein an animal learns to predict outcomes and adapt its behaviour via a conditioned response (Pavlov, 1927; Rescorla, 1988). For example, when a rat hears a tone shortly before a shock, it subsequently freezes upon hearing the tone the next time it is presented. These kind of conditioned responses, and capacities for associative learning more generally, have been widely documented across the animal kingdom, from honeybees and fruit flies to arthropods and cephalopod mollusks, and are thought to be realised neurally through mechanisms of synaptic plasticity (Brembs & Heisenberg, 2000; Buchanan & Bitterman, 1988; cf. Gallistel & Gibbon, 2010; Hawkins & Byrne, 2015; Jozet-Alves et al., 2023; see Martin et al., 2000 for a review).^1^

Despite ongoing debates about the methodological requirements for demonstrating associative learning within conditioning paradigms and the individuation of necessary and sufficient criteria for associative learning proper, its theoretical underpinnings are considered to be mature (Mackintosh, 1974; Pearce & Bouton, 2001; Shanks, 1995; cf. Gallistel, 2008; Gershman, 2025). However, disagreements have proliferated in recent decades, spurring progress not only in comparative cognition and psychology but also cognitive evolution, particularly with respect to the fundamental mechanisms of learning (Barron et al., 2023; Halina, 2022; Ginsburg & Jablonka, 2010, 2021; Schnell et al., 2021). In particular, more phylogenetically distant species that ostensibly show adaptive behaviour which, at least on its surface, appears to be consistent with capacities for associative learning, are at times excluded on account of particular criteria (Currie, 2021; Dacey, 2016; Dickinson, 2012). Other accounts argue for a more inclusive view, in which our tendency to continually shift the goal-post to exclude such organisms is evidence of our cognitive biases in biological research (Lyon et al., 2021), although some caution for a more rigorous and systematic methodology for doing so (Figdor, 2022, 2024). In order to reconcile these diverse perspectives, a more comparative approach that is sensitive to both species-specific constraints and properties of a given species’ phenotypic niche is required in order to disentangle how associative and non-associative effects contribute to adaptive behaviour, and a more careful examination of the role of ecological pressures in driving cognitive evolution (Andrews, 2020; Ginsburg & Jablonka, 2010; Halina, 2023; Shettleworth, 2010).

The standard mechanistic story defines associative effects in terms of physical mechanisms of memory, particularly through the modification of synaptic responses and long-term potentiation that associate cues and outcomes (Fanselow & Poulos, 2005; Geinisman et al., 2001; Gruart et al., 2006; Maren, 2005), which naturally precludes learning of this kind in systems without neuronal machinery. Whether that assumption holds is an empirical question—and one that perhaps requires more nuance than a yes/no dichotomy to testing associative learning (Starzack & Gray, 2021). Several traditions argue that it does not hold, though the evidence remains limited and contested (Fernando et al., 2009; Gallistel & Matzel, 2013; Gandhi et al., 2007; Hennessey et al., 1979; McConnell, 1966; Tagkopoulos et al., 2008; Walters & Byrne, 1983; cf. Loy et al., 2021).

These debates about the potential for learning in non-neuronal organisms have in fact been recurring for over a century. Initial studies were conducted—and subsequently elaborated upon—in the late 19th and mid 20th centuries purporting to demonstrate conditioned responses in “lower organisms” (Jennings, 1906). This included several unicellular ciliates, particularly the bacteria *Paramecium* (*P. caudatum* and *P. aurelia*) and marine feeder *Stentor* (*S. roeselli* and *S. coeruleus*) that show altered taxis behaviour after repeated pairings with food and “trial-and-error” like avoidance responses (Gelber, 1952, 1956; Jennings, 1904, 1906; see Gershman et al., 2021 for a review). Similar debates have ensued following reports of associative learning in plants and a constellation of other intrepid protozoa (Fernando et al., 2009; Gagliano et al., 2016; Brenner et al., 2006; Markel, 2020; see Loy et al., 2021 for a critical review). Typically, a positive test of some complex capacity is reported, before an alternative, more “low-level” account of the learning processes is proposed, maintaining a broad consensus that advocates some version of Morgan’s Canon (1894).^2^ In this case, rather than associative conditioning, many accounts instead argue that what is observed is better explained by non-associative processes like sensitisation or habituation. Drawing strong conclusions from this body of work is further complicated by the lack of adequate controls that would rule out non-associative learning effects (Loy et al., 2021). In other cases, critics have also tried and failed to replicate protocols (Markel, 2020). Overall, several methodological requirements remain unmet in the study of associative learning in non-neuronal organisms (Kehoe, 2014; Gallistel, 1990; Ginsburg & Jablonka, 2010; cf. Doan et al., 2026), but also in organisms with a comparably simple nervous system, such as the planarian flatworms *G. dorotocephala* and *S. mediterranea* (Kelso et al., 2026). There does not currently appear to be a “smoking gun” by which to quell these conceptual and methodological disputes.

*Physarum polycephalum* (*P. polycephalum*) is a particularly striking model organism by which to try and address the question of associative learning in non-neuronal organisms. Although unicellular, during the plasmodial stage of its life-cycle—in which its distributed, multinucleated, excitable membrane rhythmically shuttles and adjusts its growth processes—*Physarum* exhibits an unusually rich behavioural repertoire. Previous work has shown that *Physarum* can solve spatial tasks and adapt to complex environmental structure, habituate to a repeated aversive stimulus and recover upon its removal, and even transfer the informational effects of such processes to genetically similar individuals through cell fusion, which suggests the presence of experience-dependent plasticity to some degree (Boisseau et al., 2016; Boussard et al., 2021; Dussutour et al., 2010; Dussutour, 2021; Nakagaki, 2001; Nakagaki et al., 2000; Reid et al., 2012; Reid et al., 2016; Saigusa et al., 2008; Severs & Wang, 2025). These findings indicate that the organism’s behaviour is shaped by internal processes capable of retaining and deploying information from prior experience that are embodied across its sensorium. However, whether such effects should be explained through appeal to associative learning in the stricter sense remains unresolved.

The question of associative learning in *P. polycephalum* has been previously raised. Most notably, Shirakawa et al (2011) claimed to demonstrate associative learning through an aversive conditioning paradigm in which the organism appeared to approach an otherwise aversive thermal cue after repeated pairing with food, a pattern resembling counterconditioning. Yet subsequent discussion has highlighted the difficulty of interpreting such findings in the absence of stronger controls, including procedures that eliminate contextual confounds and other non-associative forms of learning, and a paucity of details about the protocol by which to attempt replication (Carrasco-Pujante et al., 2021; De la Fuente et al., 2019; Loy et al., 2021). In order to address these limitations, an experimental design that provides a well-defined set of hypotheses and behavioural criteria by which to operationalise associative and non-associative processes and constrain theorising, “including errors, limits, and biases so as to constrain the cognitive hypothesis space effectively” (Taylor et al., 2022, p. 3), is required. However, these approaches must also remain sensitive to the lived environment of the model organism under study, using naturalistic stimuli to elicit behavioural responses wherever possible (Sims, 2023). Perhaps most importantly, results should be interpreted cautiously and in line with their evidential strength in order to delimit further sources of bias within analyses (a problem that is particularly common in the field of comparative cognition; Shettleworth, 2010, cf. de Waal, 1999).

In the present study, we implemented a classical conditioning paradigm in which blue light (CS) was paired with a nutrient (US) across repeated training and testing cycles to examine signatures of associative learning in *P. polycephalum*. Plasmodia are sensitive to several wavelengths and generally show aversive responses to illumination, particularly in the blue range, with effects on movement and physiology (Adamatzky, 2013; Rakoczy, 1980; Starostzik & Marwan, 1995; Wohlfarth-Bottermann & Block, 1981). By contrast, nutrient sources such as oat flakes provide an essential, appetitive stimulus. Placing the food within the boundary of a projected blue light region pushes the plasmodia to cross an aversive zone to feed. We thus sought to establish: (1) whether blue light acts as an aversive or neutral stimulus, (2) whether training could induce a conditioned response in which the location of the blue light becomes associated with the presence of oat, leading to reduced avoidance or preferential migration behaviour, and (3) to establish a more reproducible methodological approach by which to identify behavioural features that may be informative to future studies. The experimental design comprised five conditions: three controls (neutral, blue light only, and oat only, all with untrained subjects), a training condition in which blue light and oat were presented together, and a testing condition in which trained subjects were exposed to blue light alone to assess conditioned responses.

Building on previous studies of *Physarum*, we quantified behavioural response using time-lapse image analysis that captured multiple dimensions of morphological and kinematic measures as well as individual trajectories. This approach enabled a broad, sensitive exploration of behavioural change while testing target, theory-driven hypotheses about what signatures, if any, are suggestive of associative learning-like effects. Our aim was not, therefore, to claim a definitive demonstration of a non-neuronal associative mechanism, but to ask whether paired exposure produces a subsequent behavioural profile that is consistent with conditioning criteria. More specifically, if blue light acquires altered behavioural significance (i.e., how the organism responds to the stimulus) through pairing with a nutrient, then plasmodia tested with blue light alone should show greater engagement with the blue light region than untrained plasmodia exposed to blue light alone. Such engagement would be expected to appear in more than one feature of behaviour, including approach probability, latency, spatial proximity and the organisation of migration.

## 2. Materials and methods

### 2.1 Organism and culture conditions

Plasmodial samples of *Physarum polycephalum* (Japanese strain) were used throughout the study. Cultures were maintained in sterile Petri dishes (SARSTEDT, 150 mm diameter, 20 mm height) on 1% agar supplemented with 5% blended oat flakes. Cultures were kept in darkness at 23°C and 80% ± 5% relative humidity. Experimental samples were excised from a common parent culture. No mating was involved, and all samples were therefore clonally derived. This approach was adopted to minimise genetic variation and other sources of variability among samples (Vogel et al., 2015). For each trial, a fresh plasmodial fragment measuring approximately 20 mm × 20 mm was excised from the culture and transferred to the centre of a fresh experimental dish containing a 1% agar base. Each fragment was used for a single experimental run within a given condition or training stage.

### 2.2 Experimental apparatus and arena

The experiment was conducted in circular Petri dishes prepared as open, unobstructed arenas, without channels or physical barriers to constrain migration. This layout was chosen to permit free expansion and migration of the plasmodium and to capture behavioural responses under minimally structured spatial conditions. The unconditioned stimulus was a moistened oat biscuit prepared from dry oat material weighing 0.5 g and measuring approximately 20 mm in diameter and 2 mm in thickness. When present, the oat biscuit was placed near the edge of the dish. The conditioned stimulus was a projected blue light area generated using a 470 nm filter (MIDOPT BP470-D40X2; projected diameter 40 mm). The filter was positioned at the edge of the Petri dish cover so that the light formed a circular blue light area on the agar surface. Dish orientation and stimulus side were randomised across trials. The projected blue light stimulus was delivered with constant distance, position, and angle across all experiments. The spatial arrangement of the stimuli is shown in Figure 1. In conditions where both stimuli were present, the oat biscuit was positioned centrally within the projected blue light area, approximately 1 cm from the dish edge. Under this arrangement, a plasmodium approaching the food source from the centre of the dish had to enter the blue light region before reaching the oat.

**Figure 1.**
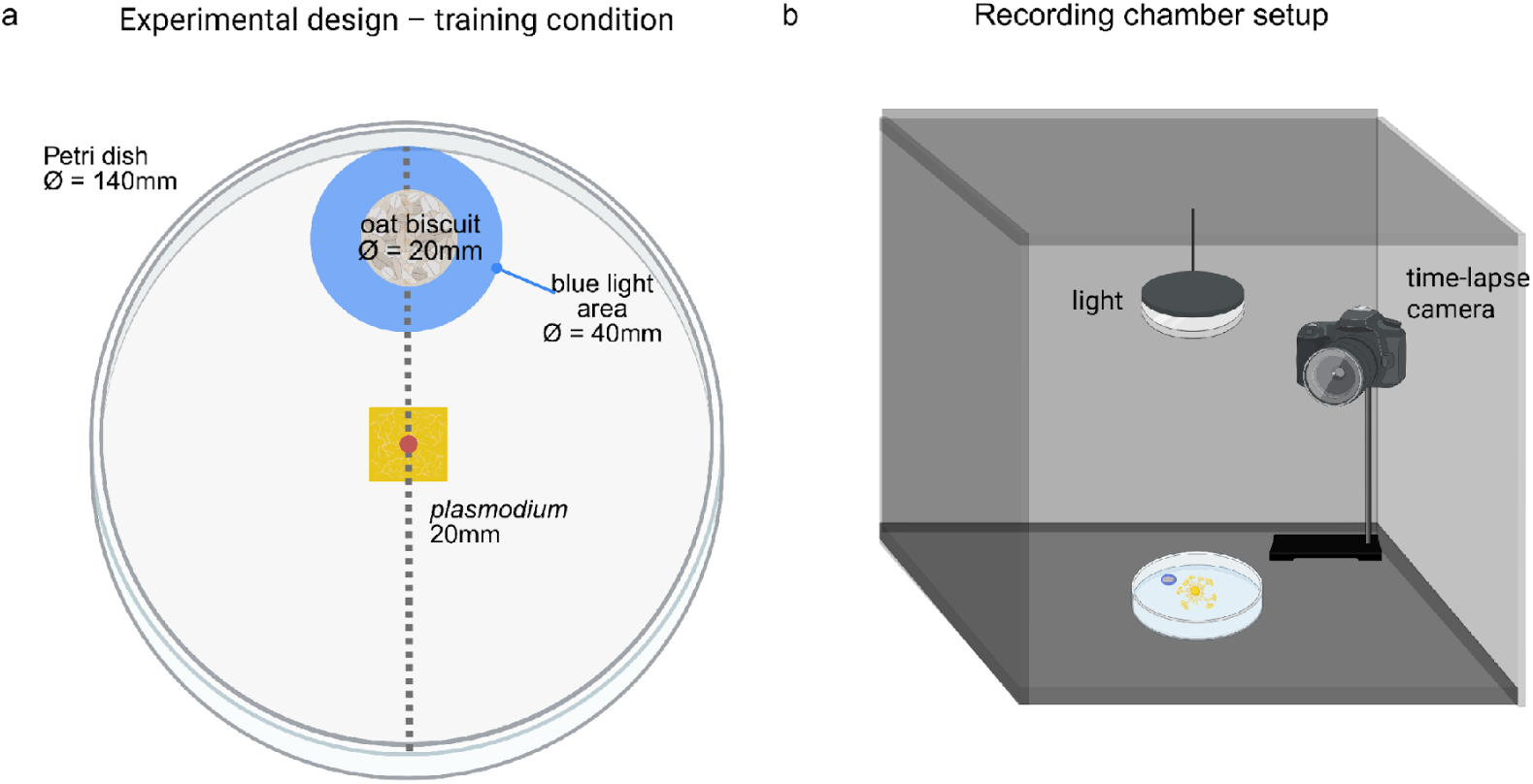
(a) Top-down view showing the position of the plasmodium sample and stimuli in the training condition, where both the unconditioned stimulus (oat) and the conditioned stimulus (blue light) are present. All other conditions use the same spatial arrangement but differ in which stimuli are present. (b) Schematic of the recording chamber, with the Petri dish placed at the centre of a light-shielded box and an overhead lamp positioned directly above the dish. Light exposure is controlled by the time-lapse camera, occurring once per minute for 1/30 s, during which an image of the plasmodium is captured.

Each trial was conducted within a light-isolated recording chamber measuring 45 × 45 × 45 cm and lined internally with a black surface to minimise external light contamination. Time-lapse images were acquired over 24 hours using an ATLI EON camera positioned above the dish. Illumination was synchronised with image acquisition and delivered as brief pulses once per minute, with each pulse lasting 1/30 s. Outside these acquisition pulses, the chamber remained dark. Ambient temperature was maintained at 23°C and relative humidity at 50% ± 5% during recording.

### 2.3 Experimental design and conditions

The study comprised five experimental conditions: neutral, blue light, oat, training, and testing (Figure 2). These conditions were designed to characterise baseline migration, responses to each stimulus presented alone, behaviour during paired stimulus exposure, and subsequent behaviour when the blue light cue was presented alone after prior pairing. The neutral, blue light, and oat conditions were untrained control conditions. In the neutral condition, the plasmodium migrated for 24 hours in the absence of both blue light and oat. In the blue light condition, the plasmodium migrated for 24 hours with the projected blue light area positioned near the edge of the dish. In the oat condition, the plasmodium migrated for 24 hours with a moistened oat biscuit placed near the dish edge.

**Figure 2.**
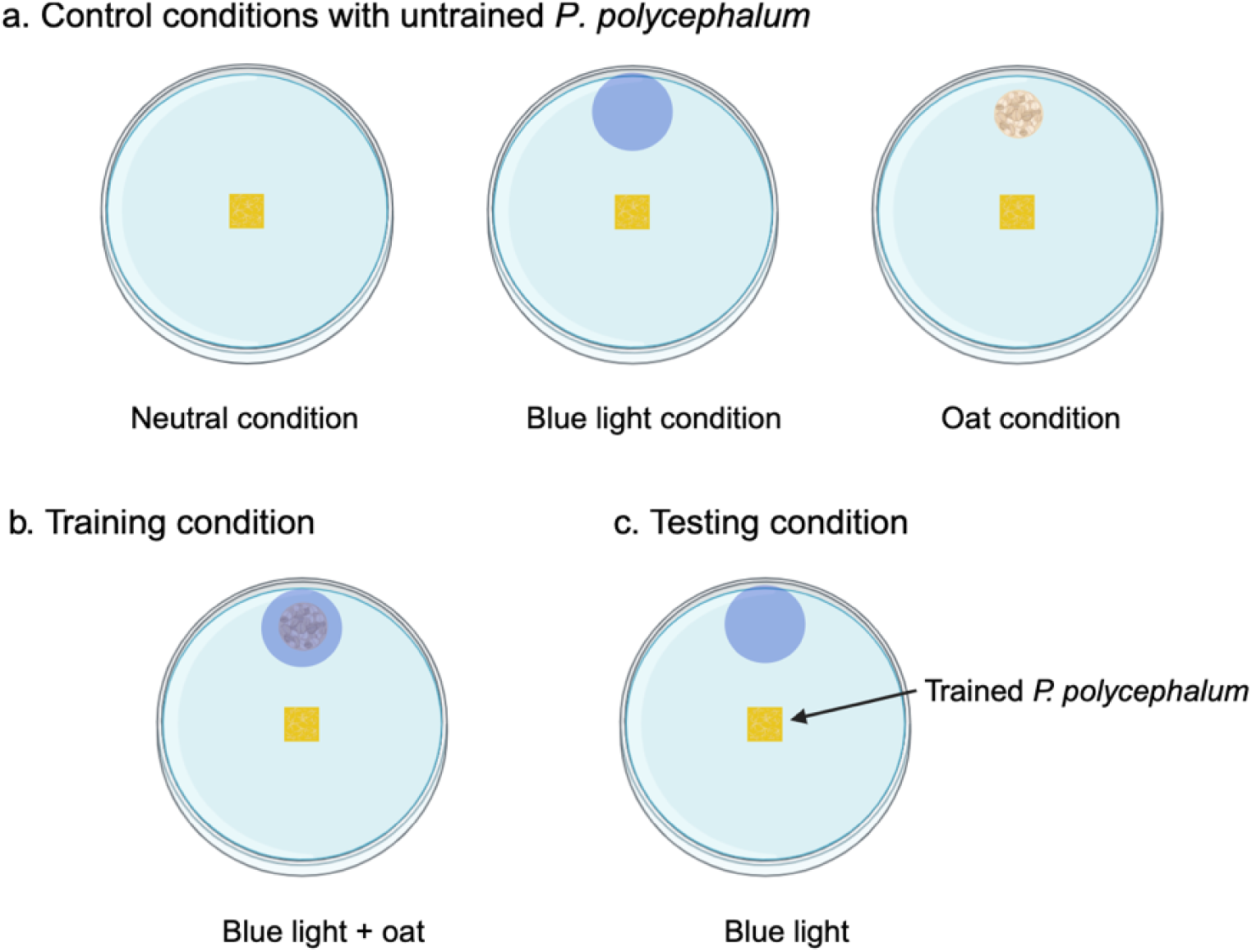
Illustration of all experimental conditions, including (a) three control conditions: neutral, blue light and oat condition; (b) training condition where oat (US) and blue light (CS) are coupled to appear, and (c) testing condition where trained *P. polycephalum* were tested by blue light only.

The training condition consisted of paired exposure to the blue light area and the oat stimulus. During training, the oat biscuit was placed directly beneath the projected blue light area so that the two stimuli co-occurred spatially throughout the 24-hour recording period. The purpose of this condition was to expose plasmodial subjects to repeated co-presentation of an aversive light cue and a food source within a consistent spatial configuration.

The testing condition was designed to assess subsequent behaviour towards the blue light area after prior paired exposure. In this condition, previously trained subjects were placed in a fresh arena and exposed to the blue light area alone, with no oat present. Behaviour in the testing condition was compared primarily with that of untrained subjects in the blue light condition. The study was designed to test whether prior paired exposure altered subsequent engagement with the blue light region at the behavioural level.

### 2.4 Training and testing procedure

Training was done in four distinct stages, designated training1, training2, training3, and training4 according to the number of completed paired-exposure cycles, following the same procedure. For training1 each subject completed a 24 hour training session and were subsequently transferred to a culture dish, only if they successfully reached the blue light area. Following the completion of training1, plasmodial subjects that successfully completed the first stage (n = 6) were collected in one culture dish and allowed to fuse and recover for 72 hours. After the recovery period, plasmodial subjects underwent a second training session (training2). Only subjects that had completed training2 were entered into the testing phase.

While training1 and training2 were interspersed by a 72h recovery period, training3 and training4 were separated from training2 and training3, respectively, by an additional seven day period during which testing took place.

Accordingly, three testing groups were analysed: testingG2, testingG3, and testingG4, corresponding respectively to subjects tested after two, three, or four completed training cycles.

During testing, the blue light stimulus was presented alone for 24 hours in a fresh arena, and behavioural responses were recorded using the same imaging setup as in the control and training conditions. In total, 84 subjects were included in the qualitative and quantitative analyses: neutral control, *n* = 10; blue light control, *n* = 10; oat control, *n* = 10; training, *n* = 24, comprising six subjects each in training1, training2, training3, and training4; and testing, *n* = 30, comprising 10 subjects each in testingG2, testingG3, and testingG4. Importantly, because *P. polycephalum* was studied as excised plasmodial subjects rather than as individually re-identifiable organisms in the animal sense, the training and testing schedule should be understood as comparing behavioural responses across clonally derived subjects with different exposure histories. The design therefore supports inference at the level of exposure-dependent behavioural change in plasmodial material, while leaving broader questions of biological individuality open.

### 2.5 Image acquisition and preprocessing

Following image acquisition, all image pre-processing, segmentation, feature extraction, statistical analysis and figure generation were performed in Python. The image analysis pipeline comprised intrinsic camera calibration, image undistortion, spatial rectification, segmentation of the plasmodial body and extraction of frame-level behavioural features.

#### 2.5.1 Camera calibration

Intrinsic camera calibration was performed for the ATLI EON time-lapse camera to estimate focal length, optical centre and lens distortion parameters. Calibration images were acquired using a planar chessboard pattern consisting of 7 × 7 squares, corresponding to 6 × 6 internal corners, with a square size of 15 mm. Chessboard corners were detected in OpenCV using *findChessboardCorners*, with adaptive thresholding and fast checking enabled (Bradski, 2000).

Object points and image points were accumulated across calibration images in which corner detection was successful, and intrinsic parameters were estimated using *calibrateCamera*. The resulting camera matrix and distortion coefficients were retained for all subsequent image pre-processing steps. Calibration images were visually inspected to confirm accurate corner detection.

#### 2.5.2 Image undistortion and spatial rectification

Each time-lapse frame was undistorted and transformed to a standardised top-down metric view. For each subject, a chessboard calibration image acquired in the same imaging configuration was used for extrinsic calibration. Four reference points corresponding to the corners of the imaging field were identified and mapped to known physical coordinates on a virtual 300 mm × 300 mm canvas, with 75 mm margins to centre the experimental arena. A homography matrix was computed with OpenCV using *findHomography*. All experimental images were first corrected for lens distortion and then warped to produce rectified orthographic images aligned to a metric grid. The rectified images were saved in TIFF format for downstream analysis.

#### 2.5.3 Definition of regions of interest and segmentation

For each subject, the Petri dish boundary and, where applicable, the stimulus region were defined in the rectified image by identifying their centres and radii. These parameters were derived from the standardised geometry of the experimental setup and verified by visual inspection. The plasmodial body was segmented from each frame using colour-based thresholding in HSV space. Threshold ranges for the yellow signal were selected empirically and adjusted at subject level where required to accommodate variation in image quality and illumination. Binary masks were refined by morphological opening and removal of small objects. The largest connected component was retained as the main plasmodial body for subsequent analysis.

#### 2.5.4 Frame-level feature extraction

Morphological features were extracted from each segmented frame using *regionprops* in scikit-image (Van der Walt et al., 2014). These included area, perimeter, eccentricity, and major and minor axis lengths. Centroid coordinates were tracked across consecutive frames to derive kinematic and trajectory-related measures, including step distance, speed, acceleration, cumulative path length, net displacement, tortuosity and mean squared displacement. Additional trajectory descriptors, including centroid path length and path entropy, were also computed. For conditions containing a defined stimulus region, goal-related features were calculated relative to that region. These included centroid distance to the stimulus area and latency to first entry. In the neutral condition, no goal area was defined, and only general morphological and kinematic features were retained. Frame-level measurements for each subject were exported as CSV files for subsequent subject-level summarisation.

### 2.6 Behavioural feature extraction

To characterise behavioural response at subject level, frame-level measurements were summarised into a multivariate behavioural profile for each plasmodial fragment. This approach was adopted because the response of *P. polycephalum* to environmental stimuli is distributed across morphology, locomotor dynamics, and spatial engagement rather than being expressed as a single discrete action. Subject-level metrics were therefore derived to capture complementary aspects of these behavioural dimensions. A total of 32 dependent variables were computed and grouped into four categories. Full mathematical definitions are provided in Supplementary Table S1.

First, *morphological variables* comprised 14 features describing organismal expansion and shape, including area, perimeter, eccentricity, and major and minor axis lengths, each summarised by mean, maximum, variability, or temporal change across the observation period. Second, *kinematic variables* comprised four features describing movement dynamics, namely mean speed, maximum speed, mean step distance, and acceleration. Third, *trajectory and goal-related variables* comprised 10 features characterising migration pattern and spatial engagement with the stimulus region. These included tortuosity, path entropy, mean squared displacement, mean centroid distance to the goal, minimum centroid distance to the goal, area at the end of migration, and time to maximum area. Fourth, *binary goal-attainment variables* comprised four outcomes indicating whether the subject entered the defined stimulus region and, if so, the latency to first entry. For the blue light and training conditions, these variables referred to the blue light area. For the oat and training conditions, equivalent variables were also computed relative to the oat region. In the neutral condition, no goal-related variables were defined.

### 2.7 Statistical analysis

All statistical analyses were conducted at subject level. Descriptive statistics are reported as means and standard deviations where appropriate. The full set of pairwise comparisons and associated descriptive summaries is provided in the Supplementary Material. Given the absence of credible prior effect size estimates for conditioning-like behavioural change in *P. polycephalum*, group sizes were determined using the resource equation method (Charan & Kantharia, 2013; Festing & Altman, 2002). This approach was used to obtain a sample size sufficient for exploratory group comparison in a system for which formal effect size estimation was not available in advance.

#### 2.7.1 Analysis of continuous behavioural variables

For each comparison, the relevant data were first assessed for distributional properties, including normality and homogeneity of variance. Normality was evaluated using the Shapiro–Wilk test, and equality of variance was assessed prior to model selection. Because distributional properties varied across features and pairwise contrasts, the choice of statistical model was made separately for each comparison rather than assumed to be uniform across all analyses. Where the data for a given comparison satisfied the assumptions of approximate normality and homoscedasticity, group differences were analysed using Bayesian parametric models, including Bayesian one-way analysis of variance for multi-group comparisons. Where these assumptions were not met, Bayesian non-parametric analogues of the Mann–Whitney U test were used for pairwise comparisons. Descriptive statistics were reported according to the observed distribution of each variable. Approximately normally distributed variables were summarised as means with standard deviations, whereas non-normal variables were summarised as medians with interquartile ranges.

Bayesian models were estimated using Markov chain Monte Carlo sampling. Posterior distributions were summarised using posterior means or medians, 94% highest density intervals, the probability of direction, and Bayes factors. Evidence strength was interpreted using the following probability-of-direction thresholds (Makowski et al., 2019; Jeffreys, 1998): values below 65% were treated as indicating no meaningful directional evidence, values from 65% to 75% as weak evidence, values from 75% to 90% as moderate evidence, values from 90% to 99% as strong evidence, and values above 99% as decisive evidence. These thresholds were used as descriptive guides for interpreting posterior support rather than as strict dichotomous decision rules. The full set of continuous behavioural features was analysed, including morphological, kinematic, and trajectory-related variables. Comparisons included both the primary condition-level contrasts among the neutral, blue light, oat, training, and testing groups, and targeted subgroup comparisons between the blue light condition and each testing subgroup separately, namely testingG2, testingG3, and testingG4.

#### 2.7.2 Analysis of binary and time-to-event outcomes

Binary outcomes relating to goal attainment, including whether a plasmodial fragment entered the defined stimulus region, were analysed separately from the continuous feature set. Group differences in the proportion of subjects reaching the blue light or oat region were evaluated using contingency-table analysis with the chi-square test of independence. For latency-based outcomes, including time to first entry into a defined stimulus region, time-to-event analysis was performed using Kaplan–Meier estimators. Differences between survival curves were assessed using the log-rank test. These analyses were used to compare the temporal dynamics of engagement with the stimulus region across experimental groups. Where appropriate, binary goal-attainment outcomes were also modelled in a Bayesian framework to estimate posterior success probabilities and between-group differences.

#### 2.7.3 Interpretation of the multivariate behavioural profile

Because behavioural response in *P. polycephalum* is not readily captured by a single discrete endpoint, inference was based on the overall pattern of differences across multiple related features rather than on any single variable in isolation. The statistical framework was therefore used to identify convergent evidence across behavioural domains, including morphology, locomotor dynamics, spatial trajectory, and engagement with the defined stimulus region. Particular interpretive weight was given to binary and latency-based measures of entry into the blue light region during testing, as these provide the most direct behavioural index of altered engagement with the cue following prior paired exposure. Morphological, kinematic, and trajectory features were treated as complementary descriptors of the broader behavioural profile.

## 3. Results

### 3.1 Qualitative overview of migration patterns

Across all experimental conditions, plasmodia exhibited active migration over the 24-hour recording period. In the neutral condition, movement was exploratory and distributed broadly across the Petri dish. In the oat and training conditions, migration was oriented towards the oat region, and subjects typically reached and occupied the food area within the recording period.

In the blue light control condition, many plasmodia did not enter the blue light region within 24 hours. Their growth pattern was typically expansive, often covering a large portion of the dish and showing broad fan-like fronts with multiple branches while not extending to the blue light region. Even when the blue light region was reached, entry generally occurred late in the recording period and was preceded by extensive lateral spread. By contrast, in the testing condition after prior paired exposure, a larger proportion of plasmodia entered the blue light region within the observation period. Relative to the blue light control, subjects in the testing condition showed more compact growth, occupied a smaller overall area, and tended to follow less laterally dispersed trajectories towards the blue light region. The final frames shown in Figure 4 illustrate this contrast between the more expansive morphology observed in the blue light control and the more compact migration pattern observed in the testing condition after two training sessions.

**Figure 4.**
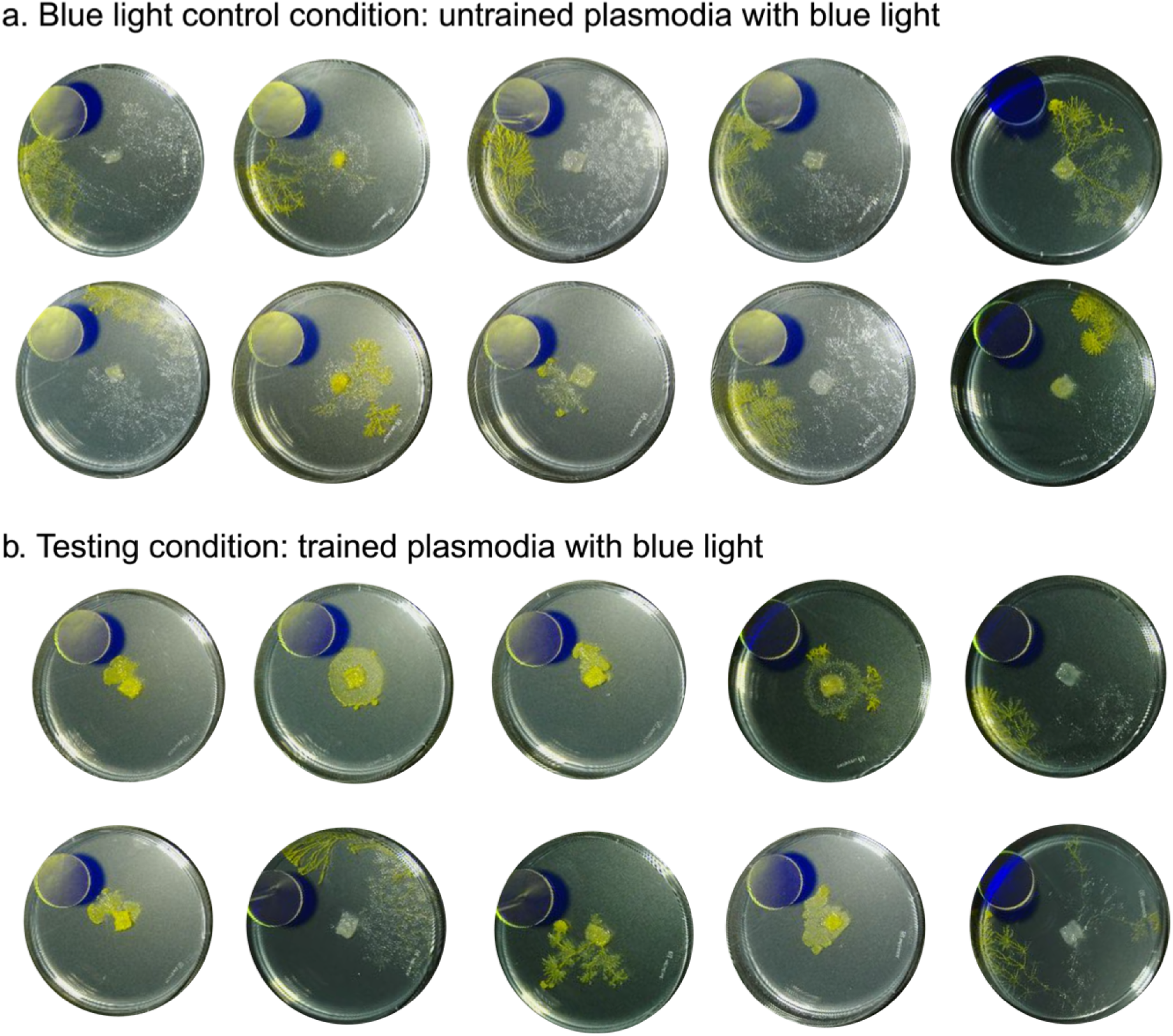
Final frames from (a) the blue light and (b) testing condition after two training sessions. Each image shows the last frame before the plasmodium either reached the blue light area or, if it did not, the last recorded frame. In the blue light control, plasmodia typically exhibited expansive, fan-like growth covering a large portion of the Petri dish, often with multiple branching fronts. When the blue light area was reached, it generally occurred late in the recording period, as indicated by extensive movement traces. In contrast, plasmodia in the testing condition displayed more compact growth, with smaller occupied area, fewer branches, and more direct trajectories towards the blue light area. A much higher proportion of subjects in the testing condition reached the blue light area, and did so with reduced lateral spread.

### 3.2 Quantitative results

The quantitative analyses encompassed a high-dimensional dataset of 32 morphological, kinematic, and trajectory-related features extracted from time-lapse recordings. *P. polycephalum* exhibits a complex and context-dependent behavioural repertoire that cannot be captured by any single measure. Migration dynamics, morphological expansion, and approach to stimuli varied across conditions, reflecting the organism’s behavioural plasticity and adaptability. To aid interpretation, Bayesian pairwise comparisons for 28 non-binary features are summarised in a heatmap (Figure 5), where colour intensity reflects the posterior probability that one condition exceeds another. Complete results and descriptive statistics for all pairwise comparisons are reported in Supplementary Table S2, 3, and targeted subgroup comparisons between the blue light control and each testing subgroup (testingG2, testingG3, testingG4) are provided in Supplementary Table S4, Figure S1.

**Figure 5.**
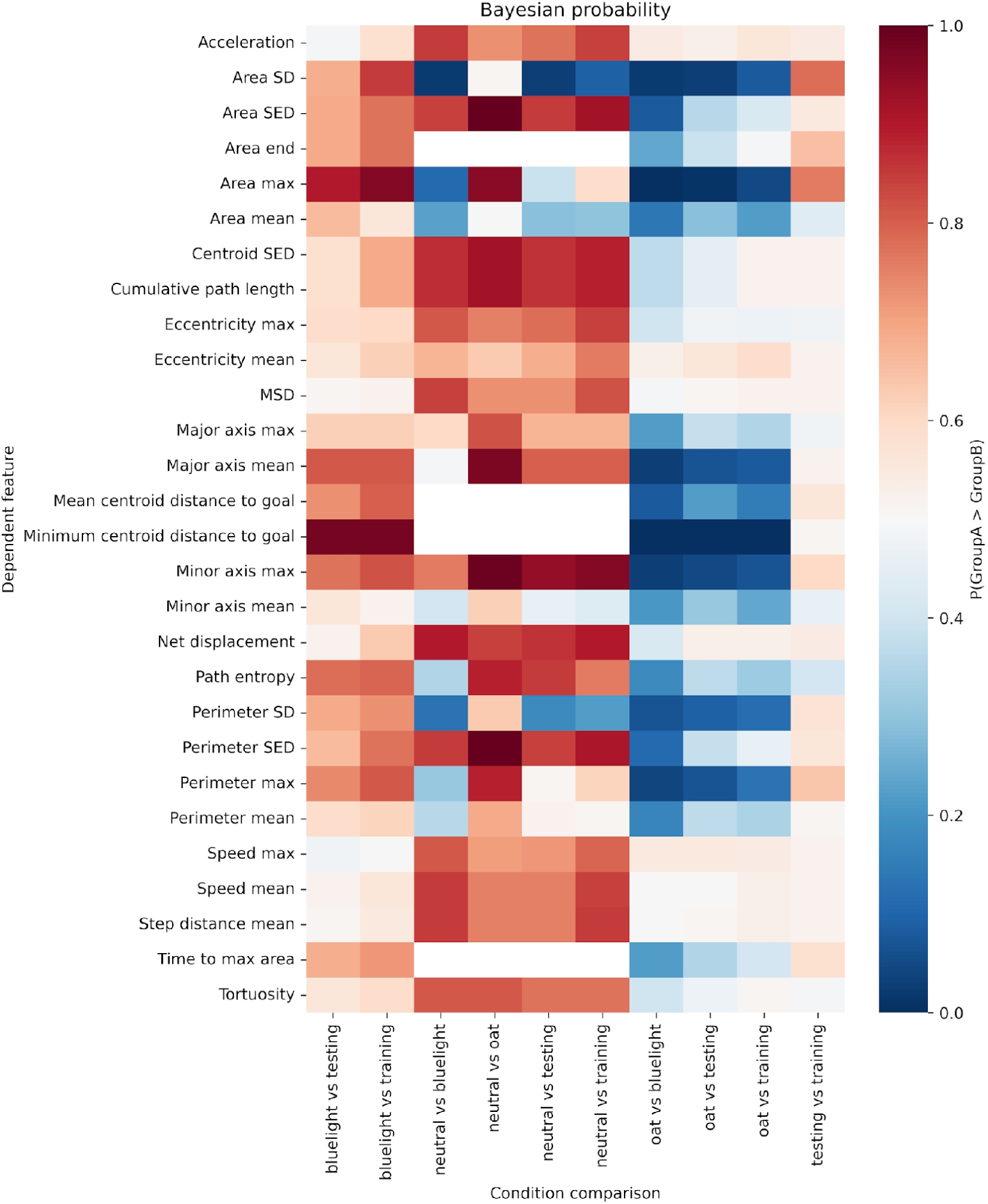
*Heatmap illustrating the posterior probability that each dependent feature is greater in Group A than Group B, for all pairwise condition comparisons.* Rows correspond to the extracted features (morphological, kinematic, and trajectory-related), and columns represent each condition comparison as group A vs. group B (e.g., blue light vs. testing, neutral vs. training, oat vs. blue light, etc.). Colour intensity indicates the strength and direction of the effect: red (values close to 1) indicates a high posterior probability that the feature is greater in the first group listed (Group A); blue (values close to 0) indicates a high probability that the feature is greater in the second group listed (Group B); white indicates little or no difference (probability near 0.5).

#### 3.2.1 Quantitative comparison of neutral and blue light conditions

Across the 28 non-binary variables, plasmodia in the neutral condition exceeded those in the blue light condition on 14 features with at least moderate Bayesian evidence (Supplementary Table S2). These differences were most evident in kinematic and trajectory-related measures, including mean speed, maximum speed, acceleration, net displacement, cumulative path length, as well as shape-related indices such as eccentricity, and tortuosity. For example, median mean speed (mm/min) was higher in the neutral condition (*Mdn* = 3.94, IQR [2.55, 7.24]) than in the blue light condition (*Mdn* = 1.27, IQR [0.72, 2.62]), and net displacement (mm) was greater for neutral (*Mdn* = 67.44, IQR [65.47, 69.53]) compared with blue light (*Mdn* = 61.57, IQR [40.16, 62.80]). In contrast, the blue light condition showed greater variability on several morphology-related and variability-related measures, including mean area, area standard deviation, perimeter, and path entropy. Relative to the neutral condition, blue light exposure was therefore associated with slower movement, reduced displacement and a broader pattern of morphological expansion. Taken together, the neutral versus blue light comparison shows that the blue light region was associated with a distinct shift in migration pattern under the present assay, affecting both movement dynamics and spatial form.

#### 3.2.2 Comparison of blue light, training, and testing conditions

Direct comparison of the training and testing conditions showed little separation across the non-binary feature set (Supplementary Table S2). Of the 28 continuous behavioural variables, 26 showed weak or no evidence of a difference between these two conditions. Moderate evidence was observed only for maximum area and area standard deviation, both of which were higher in the testing condition than in the training condition. Overall, the non-binary behavioural profiles of the training and testing groups were therefore similar. By contrast, both the training and testing conditions differed clearly from the blue light control. Across the morphology-related variables, subjects in the training and testing conditions showed lower mean area, lower maximum area, lower perimeter, shorter major axis length, and lower path entropy than subjects exposed to blue light alone. These differences indicate that migration under training and testing was associated with a more compact and less laterally expansive growth pattern than that observed in the blue light control condition.

Trajectory-related differences showed a similar pattern. Relative to the blue light control, subjects in both the training and testing conditions maintained closer proximity to the blue light region. The clearest difference was observed for minimum centroid distance to the blue light area, which was smaller after paired exposure than in the blue light control condition. Mean minimum centroid distance was 49.38 mm (SD = 6.67) after paired exposure and 55.82 mm (SD = 6.95) in the blue light control, with strong Bayesian evidence for the difference (Δ = 6.29 mm, 94% HDI [0.31, 12.42]).

The binary and latency-based outcomes provided the most direct comparison of engagement with the blue light region. A chi-square test of independence showed a significant association between condition and the probability of entering the blue light area, χ²(2, N = 64) = 19.17, *p* < .001. In the blue light control condition, 50% of subjects entered the blue light region within the 24-hour recording period, compared with 100% in the training condition and 93% in the testing condition (Figure 6a). Bayesian logistic regression showed strong to decisive evidence for higher entry probability in both the training condition (posterior mean log-odds = 2.34, SD = 0.57, 94% HDI [1.22, 3.36]) and the testing condition (posterior mean log-odds = 1.96, SD = 0.47, 94% HDI [1.09, 2.86]) than in the blue light control condition (posterior mean log-odds = 0.003, SD = 0.55, 94% HDI [−1.04, 1.04]). Model-estimated probabilities derived from the posterior means were 50.1% for blue light, 87.7% for testing, and 91.2% for training.

**Figure 6.**
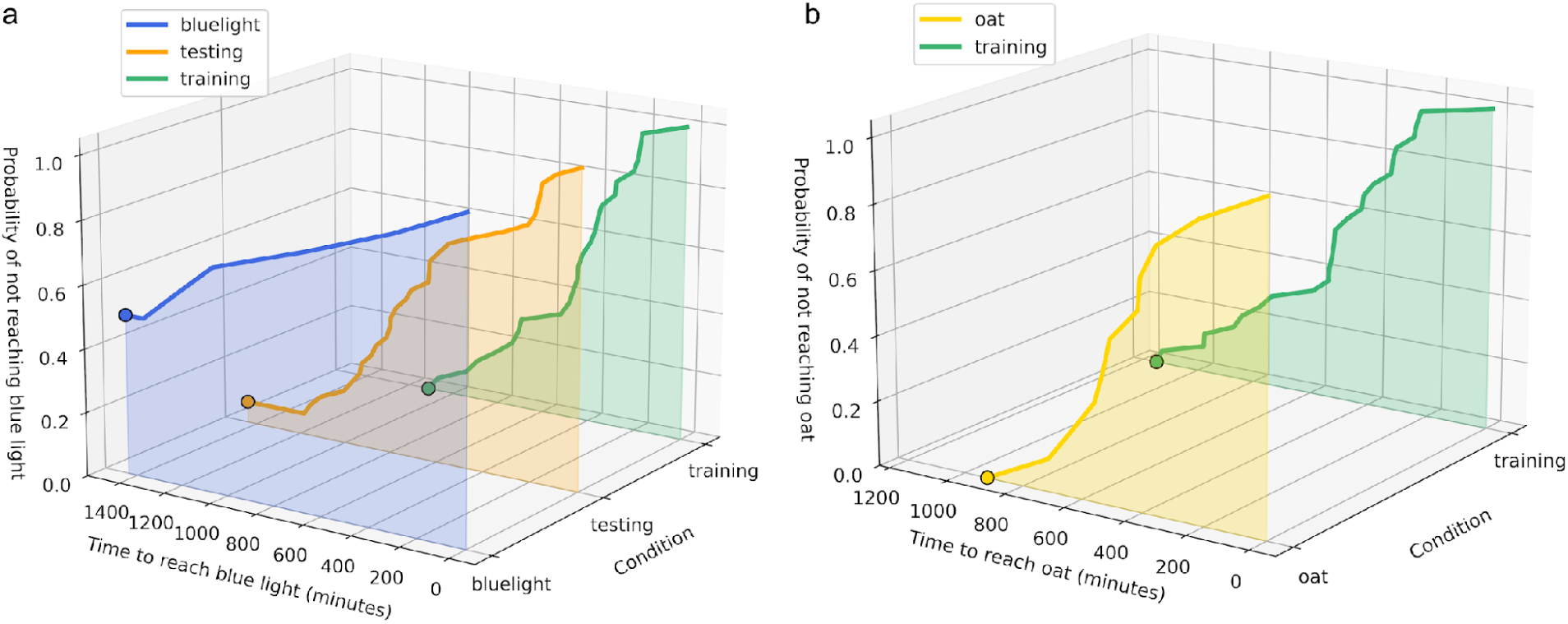
*Probability of reaching the blue light vs oat.* (a) Kaplan–Meier survival waterfall plot illustrating the probability that subjects in the blue light (blue), testing (orange), and training (green) conditions had not yet reached the blue light area over 24 hours of recording time. (b) Equivalent plot showing the probability of not reaching the oat in the oat (yellow) and training (green) conditions. Each curve represents a separate experimental condition, plotted as a function of time (minutes) to stimuli and condition. Shaded areas indicate the survival trajectory for each condition, with steeper declines reflecting faster and more consistent goal-reaching behaviour.

Latency to first entry into the blue light region also differed significantly among groups. Survival analysis showed a group effect on time to reach the blue light area, log-rank χ²(2) = 19.47, *p* < .001. The training group reached the blue light region most rapidly and consistently, followed by the testing group, whereas the blue light control group showed the slowest and least consistent entry pattern over the 24-hour period (Figure 6a).

Taken together, these comparisons show that behaviour in the training and testing conditions differed from the blue light control both in the likelihood and timing of entry into the blue light region and in the broader morphology and trajectory of migration.

#### 3.2.3 Comparison of oat and training conditions

When behavioural engagement was evaluated relative to the oat stimulus, all subjects in both the oat and training conditions reached the food source within the recording period. The difference in success rate between the two groups was therefore null, χ²(1, N = 34) = 0, *p* = 1.00. Similarly, latency to reach the oat did not differ significantly between the oat and training conditions, log-rank χ²(1) = 0.77, *p* = .379 (Figure 6b). Bayesian logistic regression likewise showed largely overlapping posterior distributions for the two groups, indicating no evidence that concurrent exposure to blue light during training reduced either the probability or the speed of reaching the oat stimulus. Despite the similarity in endpoint success and latency, the non-binary feature set indicated differences in migration pattern between the oat and training conditions. Across the 28 continuous behavioural variables, the training condition exceeded the oat condition on 13 features, most of which were morphological. In particular, maximum area was greater in the training condition (M = 1732.04 mm², SD = 521.93) than in the oat condition (M = 1355.90 mm², SD = 315.82), with strong Bayesian evidence. Additional differences included larger perimeter measures, longer major and minor axis lengths, and greater centroid distance to the blue light region in the training condition. These results indicate that although subjects in both conditions reached the oat with comparable success and timing, the broader morphology and spatial organisation of migration differed when the food source was paired with blue light.

#### 3.2.4 Comparisons among testing subgroups

Targeted subgroup comparisons between the blue light control and the individual testing groups showed that the differences observed in the overall testing condition were already present after two paired-exposure cycles. Relative to the blue light control, all three testing subgroups, testingG2, testingG3, and testingG4, showed moderate evidence for higher centroid displacement and more direct approach towards the blue light region. Differences among the testing subgroups themselves were small. Across most features, evidence for separation between testingG2, testingG3, and testingG4 was weak or absent, and there was no consistent indication of progressively stronger change with additional training cycles. The behavioural pattern observed after two pairings was therefore broadly similar to that observed after three and four pairings, with the principal divergence from the blue light control already evident at testingG2. This subgroup pattern is illustrated in Supplementary Table S4 and Figure S1.

## 4. Discussion

### 4.1 Principal findings

The present study examined whether repeated paired exposure to blue light and nutrient was associated with subsequent changes in phototactic behaviour towards an otherwise aversive stimulus. Across the assay, blue light alone was associated with a distinct pattern of migration relative to the neutral condition, characterised by slower movement, reduced displacement, and stretched morphological expansion. Against this baseline, both the training and testing conditions showed a different behavioural profile. Most notably, plasmodia previously exposed to paired blue light and oat were more likely than untrained blue light controls to enter the blue light region, and did so with shorter latency. This pattern was accompanied by differences in morphology and trajectory, including reduced lateral spread and closer proximity to the blue light area.

The clearest evidence arose from the binary and time-to-event outcomes. In the blue light control condition, half of the subjects entered the blue light region during the 24-hour observation period, whereas entry occurred in all subjects during training and in almost all subjects during testing. Likewise, latency to first entry differed significantly among groups, with the training group reaching the blue light region earliest, followed by the testing group, and the blue light control showing the slowest and least consistent entry pattern. These results provide the most direct indication that prior paired exposure was associated with altered subsequent engagement with the blue light cue.

The comparison between oat and training conditions further clarifies the behavioural pattern. All subjects in both groups reached the food source and neither success rate nor latency to oat differed significantly between them. At the same time, the broader migration profile differed, with the training condition showing a more expansive morphology. This indicates that concurrent exposure to blue light during training did not prevent subjects from reaching the nutrient source, but was associated with a different spatial organisation of growth during that process. These results indicate that paired exposure altered subsequent behaviour towards blue light while leaving the basic capacity to reach food intact.

### 4.2 Interpretation of the behavioural pattern

The behavioural pattern observed here is consistent with a conditioning modification of subsequent response to blue light. Blue light alone was associated with a characteristic baseline migration profile, whereas prior paired exposure with oat was followed by increased likelihood of entry into the blue light region, shorter entry latency, and a more compact pattern of movement towards that region. This combination of effects is compatible with the idea that the appraisal of the blue light cue had changed through repeated co-presentation with the nutrient. At the same time, the present findings should be interpreted at the behavioural level rather than as direct evidence of associative mechanism. The study was designed to examine whether a cue that is ordinarily associated with a distinct baseline response would later be engaged differently after repeated pairing with food. On that question, the results are clear enough to warrant attention. They show that prior paired exposure was associated with subsequent behavioural change under presentation of the cue alone. What they do not yet show is whether this change depends specifically on temporal contingency in the formal sense used in classical conditioning research. For that reason, it is more appropriate to describe the present findings as consistent with conditioning criteria, or as conditioning-like, than as a textbook example of associative learning.

This distinction is important in the context of debates on learning in non-neural systems. In *P. polycephalum*, as in other non-neural organisms, plasticity of the behavioural response may reflect internal physiological reconfiguration without requiring neural computation or representational mechanisms. The present results do not resolve the underlying substrate of the effect. However, they do add to a growing body of work showing that the organism’s behaviour is shaped by prior experience in ways that are not adequately described as fixed stimulus-response contingency alone. This hints at the potential novelty of these preliminary findings.

### 4.3 Alternative explanations and current limitations

A central challenge in interpreting behavioural change after repeated stimulus exposure is distinguishing associative-like effects from non-associative processes. In the present case, simple habituation does not provide an entirely satisfactory account of the observed pattern. A straightforward habituation account would predict attenuation of response to the blue light cue with repeated exposure, potentially leading to progressively reduced avoidance or increasingly indifferent migration. However, the testing subgroup analysis did not show a monotonic increase in engagement with blue light across repeated training cycles. Rather, the principal divergence from the blue light control was already evident after two paired exposures, with relatively small differences among testingG2, testingG3, and testingG4 thereafter. This pattern is not easily described as a simple cumulative habituation profile.

More generally, it is possible that bias is introduced by the training procedure. Successful training was a prerequisite, meaning that only subjects that approached the blue light and oat region were included in subsequent training and testing runs. This suggests that our results may be at least partially accounted for by a selection bias in which subjects that had a tendency to approach an aversive stimulus regardless of conditioning protocol were selected.

Equally, a simple sensitisation account is not strongly supported by the data. If repeated exposure had produced a progressively heightened aversive response to blue light, one would expect reduced entry into the blue light region or increasingly avoidant migration. The opposite pattern was observed. Relative to blue light controls, subjects in the testing condition were more likely to enter the blue light area, reached it sooner, and showed trajectories that remained closer to the blue light region. These features are difficult to reconcile with a purely sensitised response to the light cue. Additionally, in an ideal scenario, the conditioned stimulus precedes the unconditioned stimulus by a given interval, which is not guaranteed by the present paradigm. For example, perhaps chemical cues diffuse and are sensed before the blue light is encountered. In which case, approach behaviour may be triggered by soluble molecules that enter membrane proteins and an ensuing cascade of tropistic sequences, which likely occurs prior to pseudopods reaching the blue light region.

Even so, the present design does not justify strong claims that non-associative alternatives have been excluded. The most important limitation is the absence of an unpaired, random, or yoked control condition in which blue light and nutrient are experienced with matched frequency and/or the same number of presentations in the absence of predictive contingency. Without such a control, it remains possible that the behavioural change reflects some combination of repeated co-exposure, altered physiological state, stress adaptation, or other carry-over effects from the training history rather than sensitivity to the predictive relation between cue and reward. The present findings therefore support the conclusion that paired exposure was associated with a subsequent change in behaviour towards blue light, but they do not yet isolate associative contingency as the sole or definitive explanation.

### 4.4 Interpretation of the subgroup pattern

One noteworthy feature of the results is that the principal divergence from the blue light control was already evident after two paired-exposure cycles, with relatively little further separation among the testing subgroups thereafter. TestingG2, testingG3, and testingG4 all differed from the blue light control in similar directions, but the behavioural change did not increase monotonically with additional training cycles. This pattern is important because it suggests that the effect observed here is more transient or non-cumulative as opposed to a cumulative strengthening of response with repeated pairings.

Several live interpretations remain consistent at this stage. One possibility is that two paired exposures were already sufficient to induce most of the measurable behavioural change, after which additional cycles produced little further gain. Another is that the expression of the effect was constrained by the broader training schedule, which included three day recovery intervals between sessions. These intervals may have allowed partial decay or reorganisation of the behavioural change, so that additional pairings did not translate into progressively stronger performance at test. Furthermore, the training sessions (e.g., t2 and t3) were separated by an additional week, suggesting the training effect could be lost and then retrained. A further possibility is that repeated handling, transfer, or physiological fluctuation across training cycles contributed to increased variability in later groups, thereby reducing any apparent monotonic pattern. The present data do not distinguish among these possibilities, but they indicate that the relationship between exposure history and subsequent behaviour is non-linear. This non-monotonic pattern also bears on interpretation of the assay itself. In more conventional conditioning paradigms, stronger responding is often expected after additional pairings, at least within certain limits. Here, the absence of clear progressive strengthening suggests either that the relevant behavioural change was acquired rapidly and then stabilised, or that the experimental schedule interacted with the organism’s physiological dynamics in a way that shaped later expression. For this reason, the subgroup results are best interpreted as an indication that its temporal dynamics require further investigation. Nevertheless, it should not be understated that some learning/training effect is found after a single exposure and persisted for more than 3 days. In future studies, looking at the effect of time after training on behavioural change (e.g., is testing1_n1 faster at approaching than testing1_n3) could help to disambiguate these possibilities.

### 4.5 Persistence of the effect and possible mechanisms

Interestingly, the training schedule imposed long delays between pairing and test. Recall that after the first pairing, subjects that entered the blue light region were pooled in a single culture dish, where they fused and grew for 72 hours before a second pairing, and test subjects were later excised from this fused plasmodium. Any trace of pairing therefore survived cytoplasmic mixing between subjects, several days of growth and excision. Retention of conditioned responding in ciliates has been reported over intervals of 3 - 24 hours, so the persistence observed here is comparably longer. During recovery, subjects spent 72 hour on nutrient-rich agar in darkness, which amounts to prolonged exposure to the US without the CS. Such exposure degrades the contingency between CS and US, and contingency degradation suppresses conditioned responding in animals and possibly in ciliates (see Gershman et al., 2021, for a review). An associative account that tracks contingency would therefore predict some loss of the effect across recovery, whereas a pairing-induced shift in a slowly changing physiological state would predict little loss. Persistence of this length provides information about future studies on underlying mechanisms. A trace that persists across fusion and excision is potentially embodied within the cytoplasm or nuclei, which protoplasmic streaming shuttles across the plasmodium. Cell-intrinsic substrates, therefore, including RNA, DNA methylation, histone modification and enzymatic modification states maintained by continual enzyme activity, become candidate mechanisms. The causal role of these mechanisms can be examined with different methods more suitable for cellular and molecular analysis in future studies. Behaviourally, however, should the effect persist through enforced dormancy, this might suggest that a stable substrate, such as DNA methylation.

### 4.6 Broader implications for non-neural behavioural plasticity

The broader significance of the present study lies in its contribution to the experimental analysis of experience-dependent behaviour in an non-neural organism. *P. polycephalum* has already attracted attention as a model for distributed sensing, adaptive movement and problem solving, and the present findings add to that literature by showing that prior paired exposure can be associated with a later change in response to a cue presented alone. Although such a result should not be treated as definitive proof of associative mechanism, it is nevertheless relevant to ongoing efforts to understand how behavioural flexibility can arise in organisms without nervous systems. Relatedly, a particular strength of the present study is methodological. The assay combined baseline and comparator conditions with time-lapse analysis of morphology, locomotor dynamics, and spatial trajectory. This broader behavioural characterisation provides a more detailed account of how migration changed across conditions and offers a useful framework for future work. In this respect, the study extends beyond earlier reports that focused primarily on coarse approach or avoidance outcomes and provides a richer basis for evaluating behavioural changes in *P. polycephalum*.

At the same time, the findings do not require strong claims about cognition in any representational or philosophically incendiary sense. The data are compatible with the idea that prior paired exposure altered internal physiological state in a way that later affected migration under blue light presentation. Such a possibility is consistent with broader discussions of behavioural plasticity in minimal systems, where internal state reactivation or distributed physiological memory may shape later behaviour without invoking neural representations. The value of the present results therefore lies less in resolving abstract debates about cognition than in showing that experimentally tractable, experience-dependent behavioural change can be studied in a rigorous comparative framework.

## Data and code availability

The dataset and code for the project are hosted on Zenodo at: https://doi.org/10.5281/zenodo.22752964

## Footnotes

1 Importantly, associative learning refers to both a “theory-laden descriptor of both the experimental protocol and the underlying psychological/neurobiological process” (Gershman, 2025).

