## Supplemental Figure 1 for "Exposure to paired blue light and nutrient alters aversive responses in Physarum polycephalum"

Table S1. Summary of dependent features extracted from *P. polycephalum* migration trajectories.

| **Category** | **Dependent features** | **Mathematical definition** |
| --- | --- | --- |
| Mophology | area mean | Mean of segmented area (mm²) across all frames |
|  | area max | Maximum segmented area (mm²) observed over all frames |
|  | area SD | Standard deviation of segmented area (mm²) |
|  | area SED | Sum of absolute differences in area between consecutive frames |
|  | eccentricity mean | Mean eccentricity of fitted ellipse (0 = circle, 1= elongated) |
|  | eccentricity max | Maximum eccentricity observed over all frames |
|  | major axis mean | Mean length (mm) of the major axis of fitted ellipse across frames |
|  | major axis max | Maximum length (mm) of the major axis of fitted ellipse |
|  | minor axis mean | Mean length (mm) of the minor axis of fitted ellipse across frames |
|  | minor axis max | Maximum length (mm) of the minor axis of fitted ellipse |
|  | perimeter mean | Mean perimeter (mm) of the segmented body across all frames |
|  | perimeter max | Maximum perimeter (mm) observed over all frames |
|  | perimeter SD | Standard deviation of the perimeter (mm) |
|  | perimeter SED | Sum of absolute differences in perimeter between consecutive frames |
| Kinematics | acceleration | Maximum acceleration (mm/min²): largest change in speed between consecutive frames |
|  | speed mean | Mean centroid speed (mm/min): average of per-frame step distances divided by time interval |
|  | speed max | The average of the top 2% highest instantaneous centroid speeds observed during the migration period, to reduce the influence of outliers or single-frame artifacts |
|  | step distance mean | Mean Euclidean distance (mm) travelled by centroid per frame: $\frac{1}{N-1} \sum_{i=1}^{N-1} d\left( {centroid}_{i}, {centroid}_{i+1} \right)$ |
| Trajectory/Goal | area end | The area (mm²) of the slime mould at the final frame of the goal-reaching analysed migration sequence. |
|  | time to max area | The elapsed time (minutes) from the start of migration until the area of the slime mould first reaches its maximum value during goal-reaching. |
|  | mean centroid distance to goal mean | The average (mean) Euclidean distance from the slime mould’s centroid to the experimental goal (e.g., oat or blue light region) |
|  | minimum centroid distance to goal | The minimum Euclidean distance between the slime mould’s centroid and the goal, observed over the entire migration period |
|  | path entropy | Entropy (bits) of the movement angle distribution: −$\sum_{k} p_{k}\log_{2} p_{k}$ where $p_{k}$ is probability of angle in bin $k$ |
|  | tortuosity | Ratio of cumulative path length to net displacement at experiment end: $\frac{path length}{net displacement}$ |
|  | cumulative path length | Total path length (mm) traversed by the centroid: sum of all frame-to-frame centroid distances |
|  | MSD | Mean squared displacement at lag 5: $\frac{1}{N-5}\sum_{i=1}^{N-5} \left[ \left( {\Delta x}_{i,5} \right)^{2}+\left( {\Delta y}_{i,5} \right)^{2} \right]$ |
|  | net displacement | Straight-line distance (mm) between first and last centroid positions |
|  | centroid SED | Sum of Euclidean distances travelled by centroid between consecutive frames |
| Binary goal feature | reached blue light | Binary variable indicating whether the subject entered the blue light area at any point (yes/no) |
|  | time to blue light | Time (minutes) from experiment start to first entry into blue light region; censored if not reached |
|  | reached oat | Binary variable indicating whether the subject reached the oat biscuit at any point (yes/no) |
|  | time to oat | Time (minutes) from experiment start to first contact with oat; censored if not reached |

Note. Features were extracted for each subject over the full migration period and used as dependent variables in statistical analyses

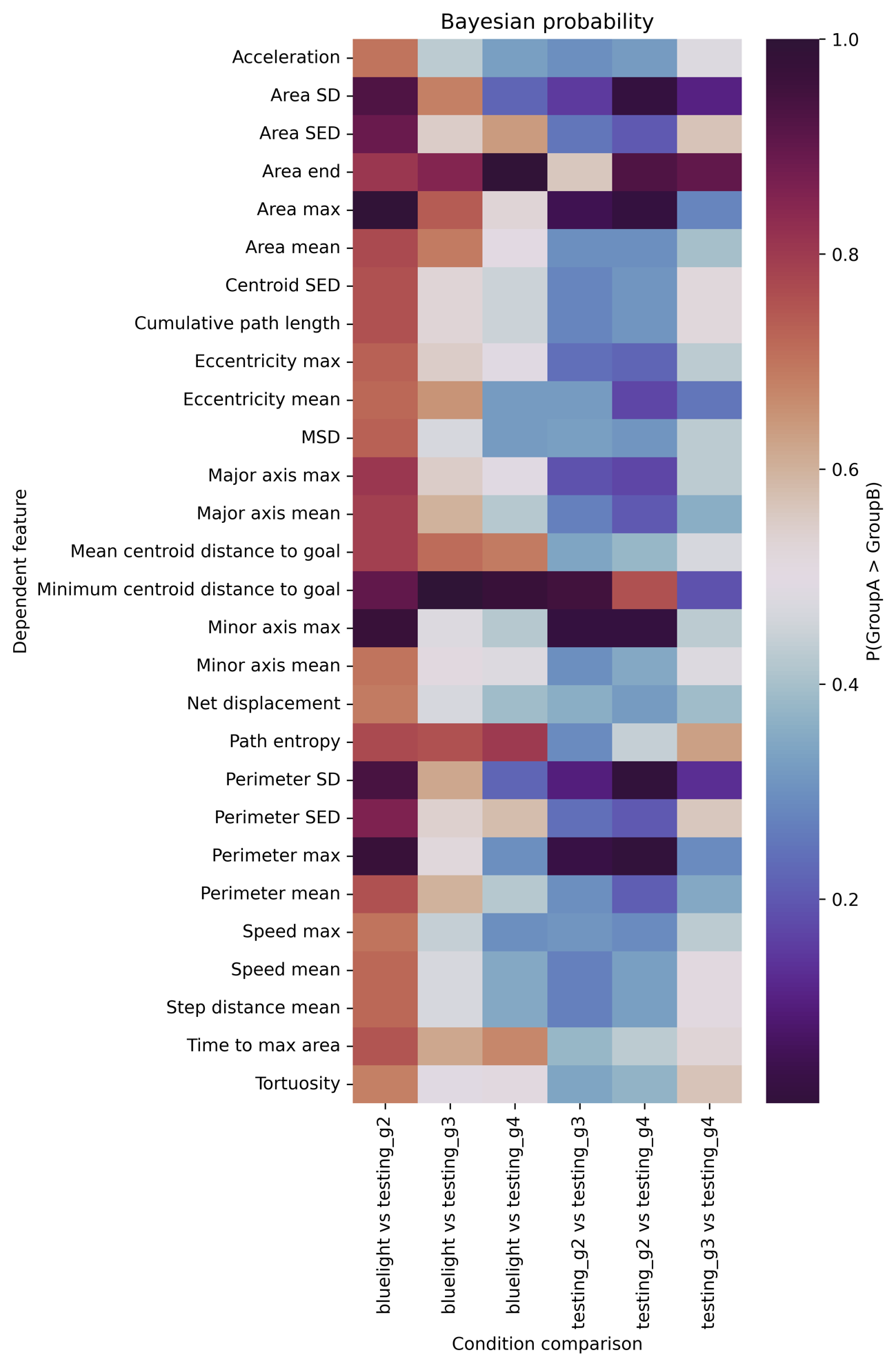

Figure S1. Heatmap illustrating the posterior probability that each dependent feature is greater in Group A than Group B, for subgroup comparisons between the blue light control and each testing subgroup (testingG2, testingG3, testingG4). Rows correspond to the extracted features (morphological, kinematic, and trajectory-related), and columns represent each condition comparison as group A vs. group B (e.g., blue light vs. testingG2, testingG3 vs. testingG4, etc.). Values close to 1 indicates a high posterior probability that the feature is greater in the first group listed (Group A); values close to 0 indicates a high probability that the feature is greater in the second group listed (Group B); probability near 0.5 indicates little or no difference.
